# Nutrient heterogeneity emerges from dynamical abiotic-biotic feedback in a spatially explicit plant-herbivore occupancy model

**DOI:** 10.64898/2026.03.03.709064

**Authors:** I. Gounand, N. Loeuille, S. Charberet, E. A. Fronhofer, E. Harvey, S. Kéfi, S.L. Leroux, C.J. Little, A. McLeod, C. Saade, F. Massol

## Abstract

Spatial heterogeneity of abiotic resources is essential for species coexistence. Ecological theory often assumes predefined heterogeneity of resources that constrains community dynamics, but the recent developments of meta-ecosystem ecology and zoogeochemistry highlight nutrient patterns could result from the interactions between the activities and movements of organisms and their abiotic environment. Here we investigate the mechanisms by which biotic-abiotic feedbacks could generate nutrient spatial heterogeneity in a simple plant-herbivore occupancy model where populations forage, recycle, and disperse in a homogenous landscape. By systematically varying organisms*’* ranges of foraging and dispersal, and recycling levels, we found that limited dispersal of plants plays a key role on the emergence of nutrient patchiness by favoring small clusters of vegetation that shape their environment through consumption and recycling. However, herbivores could also create nutrient spatial heterogeneity when large foraging and dispersal ranges, and high recycling, allow them to efficiently track plant hot spots and to increase population persistence. Unexpectedly, strong aggregation of herbivore populations did not necessarily result in nutrient clustering. Rather than via recycling, herbivores mainly affected nutrient distribution indirectly, through their top-down impact on plant distribution. When evenly spread in the landscape, herbivore populations with large foraging ranges created areas of strong herbivory pressure unfavorable to plant colonization where nutrient can accumulate.

These results can help understand the dynamical feedback between biota and abiotic resources. In a context where human activities alter both nutrient distribution and species abundances, a better understanding of this biotic-abiotic feedback will be key to anticipate the response of ecosystems to current perturbations.

## 1. Introduction

The spatial heterogeneity of abiotic resources (i.e., nutrients) has consequences for the spatial distribution and dynamics of individual organisms (e.g., Leroux et al. 2017), populations (e.g., McNaughton 1988), and communities (e.g., Harpole et al. 2011). This heterogeneity in nutrient abundance has often been construed as an external constraint and attributed to climatic and geological phenomena varying from place to place. For instance elevation and the distribution of bedrock types can influence vegetation composition (Divíšek & Chytrý, 2018), potentially cascading through the biological community by herbivore preference for certain plant (Siewert & Olofsson, 2021). Classic theoretical works have addressed the dynamical consequences of such externally imposed spatial heterogeneity in resources on biological communities. For example, MacArthur & Levins (1964) demonstrated how the spatial distribution of resources relative to movement scales affects the outcome of species competition, while optimal foraging theory showed how resource spatial distribution might influence animal movement assumed to maximize their foraging efficiency (Charnov, 1976).

However, most nutrients used by terrestrial primary producers come from biomass recycling rather than directly from bedrock or atmospheric deposition (Cleveland et al., 2013), and thus cannot be considered as an external forcing affecting communities. The recent developments of the meta-ecosystem framework (Loreau et al. 2003; Gounand et al. 2018) and of zoogeochemistry (sensu Schmitz et al. 2018) have highlighted that animals can be important contributors of nutrient cycling (Metcalfe et al., 2014), and move nutrients within and across ecosystems through local consumption, movement, and recycling processes (McInturf et al., 2019), thereby modifying the spatial distribution of nutrients. Altogether, this suggests that spatial heterogeneity of nutrients should be seen as the result of a dynamical feedback loop linking abiotic resources to biotic processes and mediated by the spatial movements of individuals.

Empirical evidence now exists regarding how animal movements affect the spatial distribution of nutrients (Gounand et al. 2018; Montagano et al. 2019). The mechanisms depend on the spatial scale through the type of movement involved (dispersal vs. foraging), and can be based on species traits affecting ecosystem processes or the movement of biotic matter (Massol et al., 2011). At a local scale, herbivores influence nutrient distribution through several processes. First, by consuming and possibly regulating plant abundance (Jia et al., 2018), they alter the local uptake of nutrients by plants and, consequently, the distribution of nutrients created by plants. Furthermore, within their home range, herbivores transport and recycle nutrients, at a scale that matches their typical movement (e.g., Schindler et Scheuerell 2002). The importance of such processes has, for instance, been highlighted in savanna ecosystems, where the recycling from large herbivores can enhance local productivity (McNaughton et al., 1997), influence plant competition (Sitters & Olde Venterink, 2021), and transport nutrients among habitat types (Veldhuis et al., 2018), such as with hippopotamus populations conveying massive amounts of nitrogen from savanna where they graze to rivers where they rest and defecate (Subalusky et al., 2015). Herbivore acceleration or deceleration of the nutrient cycle, through the balance of consumption, recycling, and loss rates (De Mazancourt et al., 1998), can even feedback to increase or decrease herbivores*’* own abundance (as proposed by Sitters & Andriuzzi, 2019). At a regional or even larger scale, species whose movement involves very large distances can couple different landscapes or regions thereby altering nutrient geographical distribution (Doughty et al., 2016; Gauthier et al., 2011). Overall, consequences of animals on nutrient fluxes can even lead to ecosystem shifts. For instance, Aleutian Islands display a grassland state where arctic foxes are absent as seabirds colonies flourish and lead to high marine-derived nutrient deposition; but where foxes are present and birds colonies controlled, the reduced nutrient flux results in a tundra ecosystem (Croll et al., 2005).

However, because plants directly consume nutrients, their role should be key in nutrient distribution, which probably does not simply reflect animal movement. Rather, the interaction between plant and animal movements (foraging, dispersal) may be key to determining the spatial structure of nutrient distribution in the landscape. Because plants are sessile organisms that consume abiotic nutrients, they are playing a direct role in nutrient patterning. This idea has been central in the understanding of vegetation patchiness, where facilitation feedbacks that lead to local infiltration of water or increased soil fixation and nutrient retention allow the formation of typical spatial structures of plant vegetation (Kéfi et al., 2007; Klausmeier, 1999). Congruently, it has been suggested that interactions with herbivores (HilleRisLambers et al., 2001) and ecosystem engineers (Tarnita et al., 2017) can largely constrain vegetation spatial distribution, and hence the underlying nutrient heterogeneity. Yet, how the spatial scales of movement, consumption and recycling of nutrients by plants, and of plants by animals, interact is still an open question. Studying the interactions between plant and herbivore spatial scales on nutrient spatial clustering using theory is one goal of the present article. Our motivation is to start with simple components to determine what patterns emerge under various contrasted scenarios in terms of movement traits (foraging, dispersal). Hopefully, the association between trait combinations and typical patterns can then be empirically tested.

While movement scales vary between taxa, they indeed also depend on the type of movement considered. Plant and herbivore species interact and mutually affect their spatial distribution, through foraging and dispersal. While herbivore foraging regulates plant populations within their home range, on a longer time scale dispersal allows plants to colonize areas with weaker herbivory pressure and more abundant nutrients. The resulting dynamical distribution of organisms should feedback on nutrient distribution through consumption and recycling. Interestingly, some of the best empirical examples of nutrient redistribution by organisms rely on recycling along animal foraging or migration movements (Croll et al. 2005; Massol et al. 2011; Beard et al. 2019). By contrast, theoretical models in community ecology mostly focus on dispersal and largely omit movements linked to foraging activities (Leibold et al., 2004; Levins, 1969). We therefore lack a clear understanding of how these two types of movement (foraging and dispersal) interact to shape the distributions of plants and herbivores. Available data suggests that dispersal and foraging spatial scales could vastly differ within a given species (Straus et al., 2024). A better understanding of the interactions between these two scales of movement is another focus of the present work.

Contrary to the classic perspective which assumes that spatial resource heterogeneity is imposed by the abiotic environment, here we ask in which conditions nutrient patchiness could emerge from organism interactions, movements, and recycling in an otherwise homogenous abiotic environment. To tackle this question, we developed a spatially explicit model that accounts for both abiotic resource dynamics (hereafter *“*nutrients*”*) and the ecological dynamics of plant and herbivore populations (each consisting in a single species). Using this framework, we investigate (1) how the scales of dispersal of herbivores and plants affect nutrient spatial heterogeneity; (2) the effect of herbivore foraging range on this distribution; (3) how nutrient patchiness depends on the level of recycling. Our results highlight how limited dispersal of plants plays a key role on the emergence of nutrient patchiness by favoring small clusters of vegetation that shape their environment through consumption and recycling. Herbivores, however, can also generate nutrient patches, especially when recycling is high and foraging range sufficiently large. We discuss the mechanisms underlying these results and their relevance to empirical systems in the light of the available evidence.

## 2. Model description

We model the meta-ecosystem dynamics of one abiotic resource and two trophic levels, observed at the scale of populations on a landscape (see Table 1 for a summary of model scale parameters). The three central variables are local stocks of an abiotic resource (hereafter nutrient), populations of primary producers (hereafter plants), and populations of herbivores (see Table 2 for a summary of variables). Similar to Holt (2002) or Calcagno et al. (2011), the model focuses on plant and herbivore occupancy and the associated colonization-extinction dynamics, and thus omits their local demographics. More details on the specifics of the model are given in Supplementary Material, Appendix 1.

**Table 1.**
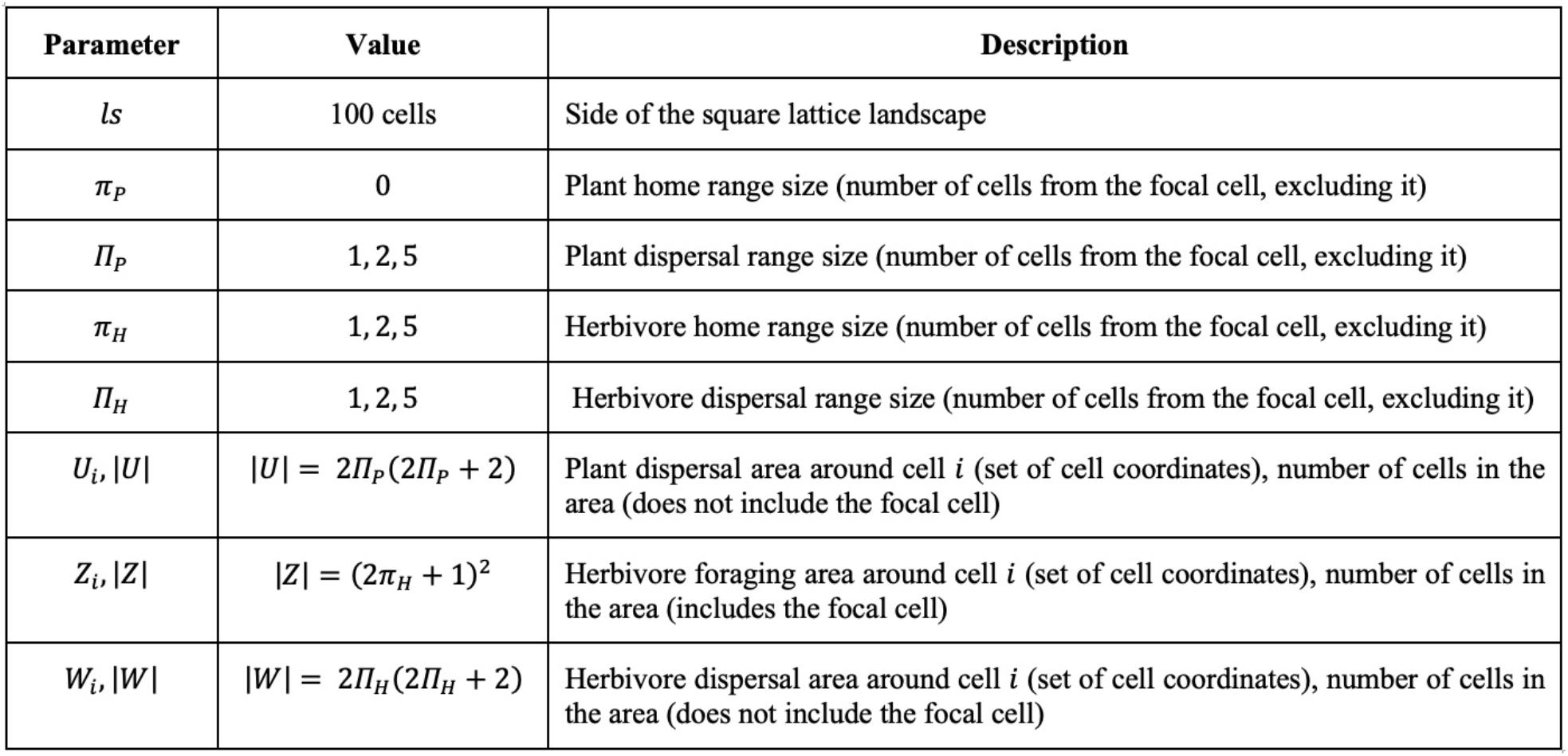
Scale and area parameters.

| Parameter | Value | Description |
| --- | --- | --- |
| $ls$ | 100 cells | Side of the square lattice landscape |
| $\pi_P$ | 0 | Plant home range size (number of cells from the focal cell, excluding it) |
| $\Pi_P$ | 1, 2, 5 | Plant dispersal range size (number of cells from the focal cell, excluding it) |
| $\pi_H$ | 1, 2, 5 | Herbivore home range size (number of cells from the focal cell, excluding it) |
| $\Pi_H$ | 1, 2, 5 | Herbivore dispersal range size (number of cells from the focal cell, excluding it) |
| $U_i, \vert U\vert $ | $\vert U\vert = 2\Pi_P(2\Pi_P + 2)$ | Plant dispersal area around cell $i$ (set of cell coordinates), number of cells in the area (does not include the focal cell) |
| $Z_i, \vert Z\vert $ | $\vert Z\vert = (2\pi_H + 1)^2$ | Herbivore foraging area around cell $i$ (set of cell coordinates), number of cells in the area (includes the focal cell) |
| $W_i, \vert W\vert $ | $\vert W\vert = 2\Pi_H(2\Pi_H + 2)$ | Herbivore dispersal area around cell $i$ (set of cell coordinates), number of cells in the area (does not include the focal cell) |

**Table 2.** Main model variables (*R*_*i*_, *P*_*i*_, *H*_*i*_) and derived variables used in the model.

| Variable | Value | Description |
| --- | --- | --- |
| $P_i, H_i$ | 0 or 1 | Plant and herbivore population presence in cell $i$ |
| $N_i$ | continuous values | Nutrient level in cell $i$ |
| $\bar{P}_{disp,i}$ | $\frac{1}{\vert U\vert } \sum_{k \in U_i} P_k$ | Average number of existing plant populations in cells within the plant's dispersal area $U_i$ around cell $i$ |
| $\bar{H}_{disp,i}$ | $\frac{1}{\vert W\vert } \sum_{k \in W_i} H_k$ | Average number of existing herbivore populations in cells within the herbivore's dispersal area $W_i$ around cell $i$ |
| $\bar{H}_{for,i}$ | $\frac{1}{\vert Z\vert } \sum_{k \in Z_i} H_k$ | Average number of existing herbivore populations in cells within the herbivore's foraging area $Z_i$ around cell $i$ |
| $\lambda_i$ | $\frac{1}{\vert Z\vert } \sum_{k \in Z_i} \frac{P_k}{\bar{H}_{for,k}}$ | Food level for herbivores in cell $i$ |

### The meta-ecosystem and biotic scales

The landscape consists of a square lattice of 100 × 100 cells, where each grid cell contains a quantitative level of nutrient, and may be occupied by a population of plants, a population of herbivores, or both (Fig. 1). A single cell corresponds, by definition, to the scale of nutrient uptake (or *“*foraging area*”*) of a plant population (i.e. the area the plant population affects through nutrient uptake). The size of the foraging area of herbivore populations is one of the main model parameters (Table 1). It is assumed to be a square, parameterized by its foraging range π_*H*_ (i.e. the maximum number of cells between the position of a herbivore population occupies and the ones it can affects through consumption), and can vary between a 3 × 3 square (π_*H*_ = 1) and a 11 × 11 square (π_*H*_ = 5). Plants and herbivores are also characterized by a second spatial scale, i.e. their dispersal ranges (*Π*_*p*_ for plants and *Π*_*H*_ for herbivores), which modulates where colonization can take place.

**Figure 1.**
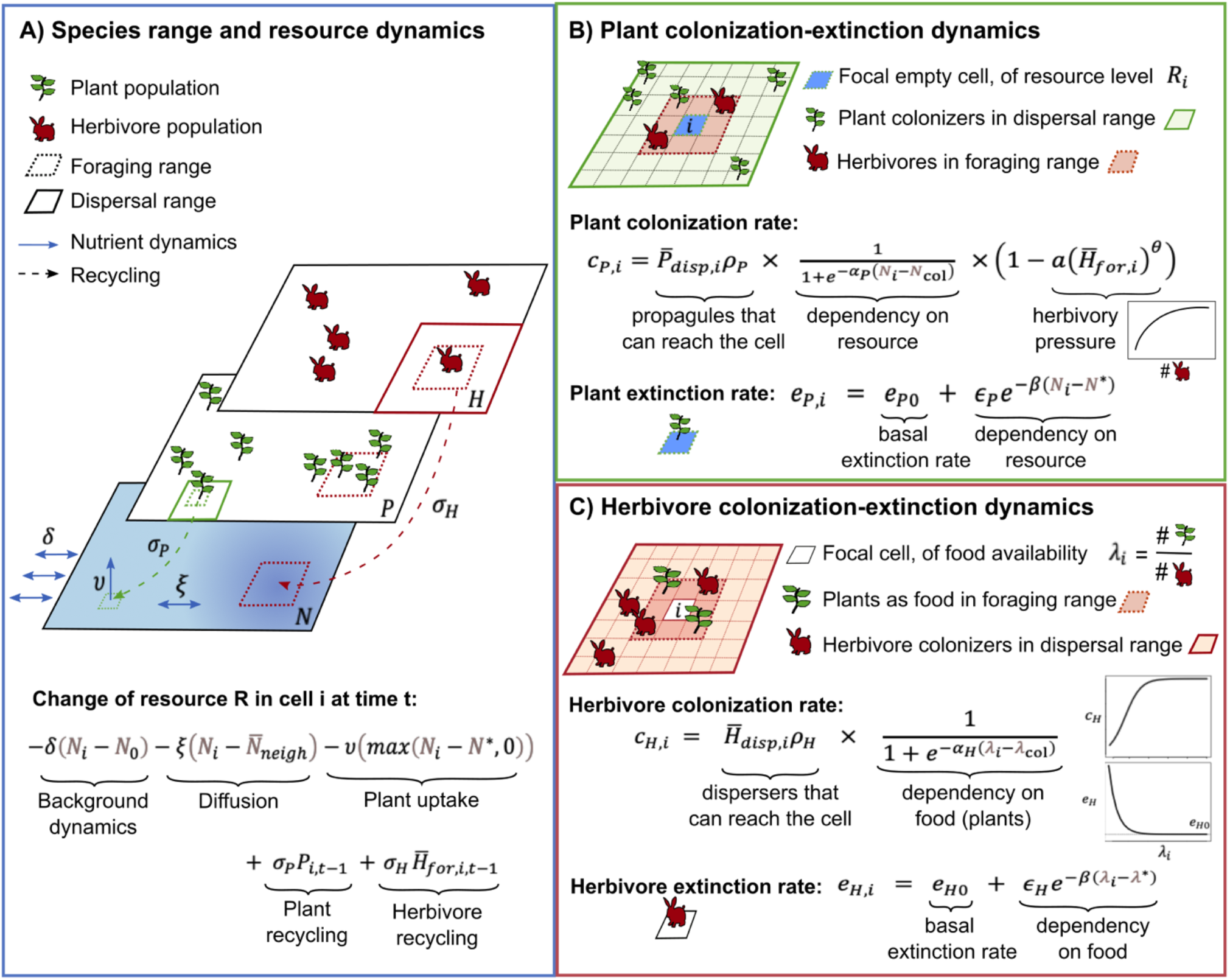
Synthetic view of the model.

### Trophic interactions and recycling

Within their respective foraging areas, plants deplete the nutrient level while herbivores consume plants and thus negatively affect the colonization rate of plant populations (Fig. 1b-c). Because herbivore populations depend on the presence of plant populations, their own colonization and extinction rates are affected by the number of plant populations occurring in their foraging area. At the same time, both plants and herbivores can positively affect nutrient levels within their respective foraging areas through recycling of detritus (Fig. 1a).

### Nutrient dynamics

In a given cell, the nutrient level (i) tends to equilibrate toward a background level *N*_0_ (e.g. through atmospheric deposition or rock weathering) following chemostat dynamics, (ii) diffuses into its eight neighboring cells, while also receiving diffusion inflows from these neighbors, (iii) is depleted by plants if there is a local plant population, and is partially replenished by the recycling of detritus of (iv) plant and (v) herbivore populations that forage on this cell (Fig. 1a). Nutrient dynamics in cell *i* is modeled in discrete time and each of these five processes contributes to its local change between time *t* and *t* + *τ* as:

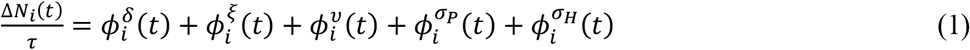

where 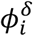 stands for the chemostat effect, 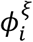, for the diffusion of nutrients from neighboring cells, 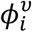, for the uptake of nutrients by the local plant population, 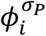 and 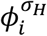, for the recycling of plant or herbivore detritus, respectively. The superscripts correspond to the scaling factors of each process (see Fig.1a, Appendix 1 for details, and Table 3 for parameter definitions). Nutrient uptake is modeled as a second chemostat, with a target resource level *N*^∗^ set by the plant (i.e. resources decrease proportionally to the difference between current resource level and *N*^∗^, similarly to Tilman 1980). Recycling is implemented as a time-lagged positive term in cells harboring a plant population 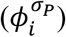 or contained within the foraging area of any number of herbivore populations 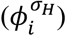. The presence of plant or herbivore populations increases the quantity of available nutrients in the next time step evenly over their respective foraging areas (i.e., local effect is total effect divided by the number of cells in foraging area).

**Table 3.** Model parameters pertaining to nutrient, plant and herbivore dynamics.

| Parameter | Dimension | Value | Description |
| --- | --- | --- | --- |
| $N_0$ | $M \cdot L^{-2}$ | 10 | Background nutrient level |
| $N_{col}$ | $M \cdot L^{-2}$ | 4 | Critical nutrient level determining the inflexion of the logistic curve relating plant colonization rate to nutrient level |
| $N^*$ | $M \cdot L^{-2}$ | 0 | Minimal nutrient level imposed by plant uptake |
| $\delta$ | $T^{-1}$ | 0.1 | Chemostat rate (input from a background nutrient pool) |
| $\xi$ | $T^{-1}$ | 0.1 | Diffusion rate of nutrient between neighboring cells |
| $v$ | $T^{-1}$ | 0.5 | Depletion rate of nutrient by plants |
| $\sigma_P$ | $T^{-1}$ | 0, 1, 2 | Recycling rate for plant detritus |
| $\sigma_H$ | $T^{-1}$ | 0, 1, 2 | Recycling rate for herbivore detritus |
| $\rho_P, \rho_H$ | $T^{-1}$ | 1.6, 2.4, 3.5, 3.6; 6, 15 | Plant and herbivore rates of propagule or offspring production |
| $\alpha_P, \alpha_H$ | ∅ | 0.75; 1 | Parameters regulating the steepness of the logistic function which relates colonization rates to nutrient level for plants and herbivores ($R_i$ for plants and $\lambda_i$ for herbivores) |
| $a, \theta$ | ∅ | 0.9; 0.5 | Herbivore pressure coefficients (strength and shape of the relationship describing this pressure, which affects plant colonization rate) |
| $\lambda^*$ | ∅ | 1 | Critical food level for herbivores (beyond which extinction rate increases dramatically) |
| $\lambda_{col}$ | ∅ | 2 | Critical food level determining the inflexion of the logistic curve relating herbivore colonization rate to food availability |
| $e_{P0}, e_{H0}$ | $T^{-1}$ | 0.4, 0.9; 1, 1.5 | Plant and herbivore basal extinction rates |
| $\beta_P, \beta_H$ | ∅ | 0.5; 1 | Coefficients regulating the steepness of the inverse exponential function relating extinction rates and nutrient levels for plants and herbivores ($R_i$ for plants and $\lambda_i$ for herbivores) |
| $\epsilon_P, \epsilon_H$ | $T^{-1}$ | 3; 3 | Scaling of extinction rates with the level of nutrients |

### Plant colonization-extinction dynamics

Plant populations may colonize cells devoid of plant populations within their dispersal range, or go extinct (Fig. 1b). Here we assume that nutrients determine both colonization and extinction, while herbivory is only affecting the colonization process. While effects of herbivory on plant colonization are often negative due to the preferential grazing of colonizing seedlings (e.g. Chabrerie et al., 2019; Schaffner et al., 2011), effects on extinction are less clear as herbivores may even reduce extinction (when relaxing competition or locally recycling nutrients) or increase it (Olff et Ritchie 1998). An existing plant population in cell *i* goes extinct at rate *e*_*P,i*_ which depends on the local stock of nutrients *N*_*i*_ through the following equation:

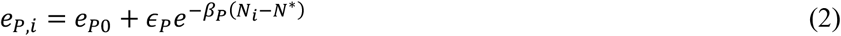

where *e*_*p0*_ is the basal extinction rate of plants, *ϵ*_*p*_ scales nutrient effects on extinction rates, *β*_*p*_ is the rate at which extinction rate decreases with the level of nutrients and *N*^∗^ is the minimum level of resources imposed by plant uptake (see Table 3 for a summary of parameters involved in plant, herbivore and nutrient dynamics). In practice, we set *N*^∗^ = 0.

A cell devoid of plants can be colonized by other plant populations within its dispersal area (i.e. all the cells present in the (2*Π*_*p*_ + 1) × (2*Π*_*p*_ + 1) square area around the focal cell). The rate *c*_*p,i*_ at which colonization occurs in a focal cell *i* is given as:

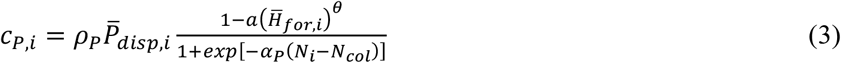

where *ρ*_*p*_ is the rate of propagule production towards each of the cells in its dispersal area by a single plant population, already discounting possible costs of dispersal, 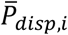 is the average occupancy of plants in the dispersal area around the focal cell, *α*_*p*_ controls the rate at which colonization rate increases logistically with nutrient level *N*_*i*_ (higher nutrient favoring colonization; e.g., Holdredge et al., 2010), *N*_*col*_ is the critical nutrient level at which the effect of a slight increase in nutrient level on colonization rate is the largest, 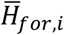 is the average occupancy of herbivores in the herbivore foraging area around the focal cell (i.e. cells are more difficult to colonize if herbivores would consume all young plants), and *a* and *θ* are parameters controlling the strength and shape of herbivores*’* negative effect on plant colonization.

### Herbivore colonization-extinction dynamics

We assume that both colonization and extinction of herbivores depend on available plants within their foraging range, accounting for competition among herbivores (Fig. 1c). An existing herbivore population in cell *i* goes extinct at rate *e*_*H,i*_, which depends on plant and herbivore occupancies in the foraging area around the focal cell *i*:

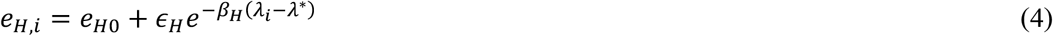

where *e*_*H0*_ is the basal herbivore extinction rate, *ϵ*_*H*_ scales the effect of the amount of available food on extinction rates, *β*_*H*_ controls the steepness of the decrease in extinction rate with the level of available food, *λ*_*i*_ is the amount of available food (see below), and *λ*^∗^ is a critical amount of food under which extinction increases exponentially (e.g. Koh et al. 2004). The amount of available food in cell *i* is defined as:

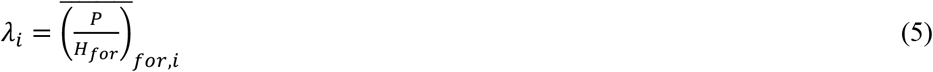

i.e. is equal to the ratio of plant populations available to herbivore populations foraging in this cell, averaged over the foraging area *Z*_*i*_ centered on cell *i*.

A cell devoid of herbivores can be colonized by other herbivore populations within its dispersal area (i.e. all the cells present in the (2*Π*_*H*_ + 1) × (2*Π*_*H*_ + 1) square area around the focal cell). The rate *c*_*H,i*_ at which colonization occurs in focal cell *i* is given as:

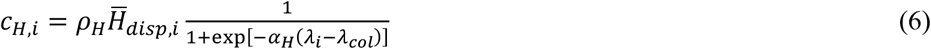

where *ρ*_*H*_ is the rate of offspring production by a single herbivore population, already discounting possible costs of dispersal, 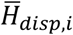 is the average occupancy of herbivores in the dispersal area around the focal cell, *α*_*H*_ is a parameter for the rate at which colonization rate increases logistically with food level *λ*_*i*_ (i.e. underexploited plants facilitate herbivore colonization), and *λ*_*col*_ is a critical food level controlling the shape of this logistic function.

## 3. Simulation and analyses

### Simulation procedure

Colonization and extinction events are modeled as Poisson processes while nutrient dynamics are ruled by a cellular automaton in discrete time. Instead of resorting to the classic Gillespie algorithm to simulate Poisson processes, we used the tau-leaping algorithm proposed by Gillespie (2001) with a constant time step, *τ*, to easily coordinate discrete biotic events with discrete-time nutrient dynamics. The model is simulated for 3000 timesteps and the value of *τ* is always equal to 0.1 (i.e. the duration of a simulation is 300 time units, a sufficient duration to reach occupancy steady states). We recorded the final landscapes at the last time point, as well as the temporal dynamics of mean nutrient values and P and H occupancies. Because the effect of organisms on nutrients increases with the time populations stays at the same place, we also computed the lifespan of plant and herbivore populations, by averaging the number of timesteps each population stays in a cell from colonization to extinction, for all colonization events occurring during the simulation. Simulations are run with or without herbivores. Simulations are always initialized with all *N*_*i*_ equal to the background level *N*_*0*_, and all cells occupied by a plant population. Simulations assume reflecting boundary conditions. To avoid possible edge effects, we discard a 15-cell boundary from the landscapes for analyses of model outputs. For scenarios involving herbivores, we randomly set herbivore populations in 20% of the cells at the onset of each simulation.

### Simulation design

To study how the scales of species movement affect nutrient spatial distribution, we systematically vary plant dispersal, herbivore dispersal, and herbivore foraging ranges, which take the values 1, 2, and 5 (squares with sides of 3, 5 and 11 cells respectively). We also varied recycling rates to assess the interactive effect of recycling with organisms*’* movement scales, considering scenarios without, or with moderate or high recycling (*σ*_*p*_ = *σ*_*H*_ = 0, *or* 1, *or* 2, respectively). Finally, we varied the parameters modulating colonization-extinction dynamics of plant and herbivore populations (*ρ*_*p*_, *e*_*p0*_, *ρ*_*H*_, *e*_*H0*_), to assess the robustness of our results for our focal questions. That is, we considered fast or slow dynamics of plants and herbivores, and dynamics leading to frequent or rare plant populations (see Table 4). We also varied *θ*, the parameter shaping herbivory pressure on plant colonization to have scenarios with more top-down (*θ* = 0.5) or bottom-up (*θ* = 1) regulation (Fig. S1), but we only show the results for *θ* = 0.5. We ran the full factorial design of these parameter variations, both with and without herbivores, and did 25 replicates for each of the 1944 scenario combinations, ending up with 48600 simulations.

**Table 4.** Scenarios corresponding to parameters variation.

| Scenario | Modality | Parameters |
| --- | --- | --- |
| Plant dispersal variation | Local dispersal | $\Pi_P = 1$ |
| | Intermediate dispersal | $\Pi_P = 2$ |
| | Global dispersal | $\Pi_P = 5$ |
| Herbivore dispersal variation | Local dispersal | $\Pi_H = 1$ |
| | Intermediate dispersal | $\Pi_H = 2$ |
| | Global dispersal | $\Pi_H = 5$ |
| Herbivore foraging variation | Local foraging | $\pi_H = 1$ |
| | Intermediate foraging | $\pi_H = 2$ |
| | Global foraging | $\pi_H = 5$ |
| Recycling variation | No recycling | $\sigma_P = \sigma_H = 0$ |
| | Moderate recycling | $\sigma_P = \sigma_H = 1$ |
| | High recycling | $\sigma_P = \sigma_H = 2$ |
| Herbivore presence | With | *no H initially present* |
|  | Without | *20% of cells occupied by H initially* |
| Plant dynamics | Fast | $\rho_P = 3.6, e_{P0} = 0.9$ |
| | Slow | $\rho_P = 1.6, e_{P0} = 0.4$ |
| | Frequent | $\rho_P = 2.4, e_{P0} = 0.4$ |
| | Rare | $\rho_P = 3.5, e_{P0} = 1.5$ |
| Herbivore dynamics | Fast | $\rho_H = 15, e_{H0} = 1$ |
| | Slow | $\rho_H = 6, e_{H0} = 0.4$ |
| Herbivory pressure | More top-down | $\theta = 0.5$ |
| | More bottom-up | $\theta = 1$ |

### Analysis of simulation outputs

At the landscape scale, we report mean nutrient level and plant and herbivore population occupancies. Spatial aggregation of nutrients, plants, and herbivores were assessed using Moran indices, computed with the R-package *‘*spdep*’*. We calculated a number of additional spatial and fragmentation statistics (see Supplementary Materials, Appendix 1), and these statistics were strongly correlated with Moran indices (Fig S2). Moran indices were normalized using z-scoring, i.e. removing the expected value obtained from 1000 independent randomizations of nutrient levels, plant or herbivore occurrences in the final landscapes and dividing this centered value by the standard deviation of the same statistic over the 1000 randomized landscapes. Thus, the z-score evaluates the observed spatial structure while controlling for variations in occupancy among scenarios. Figure S3 show examples of nutrient landscapes with extreme and median Z-scores obtained over our simulations. All analyses were run using R v.4.5.2.

## 4. Results

### 4.1. Contrasting bottom-up and top-down effects of plant and herbivore ranges on nutrient levels and species occupancies

Increasing plant dispersal range *Π*_*p*_ results in higher plant occupancy as plants can access empty cells more easily. This negatively affects the nutrient level (N), which plants (P) consumes, but increases herbivore (H) occupancy through bottom-up control (Figs. 2A, 2D, 2G). By contrast, increasing herbivore dispersal range *Π*_*H*_ allows the herbivores to better track plants and increase their occupancy, thereby slightly decreasing plant occupancy for the benefit of mean nutrient level N, relaxed from plant uptake (Figs. 2B, 2E, 2H). This top-down control of plant occupancy is accentuated when increasing herbivore foraging range, π_*H*_, as can be seen with stronger negative correlations between adjacent trophic levels (Figs. 2C, 2F, 2I). Adding recycling simply increases mean nutrient level N and both P and H occupancies by retaining nutrients in the system without affecting the patterns (Fig. 2 open versus solid points).

**Figure 2.**
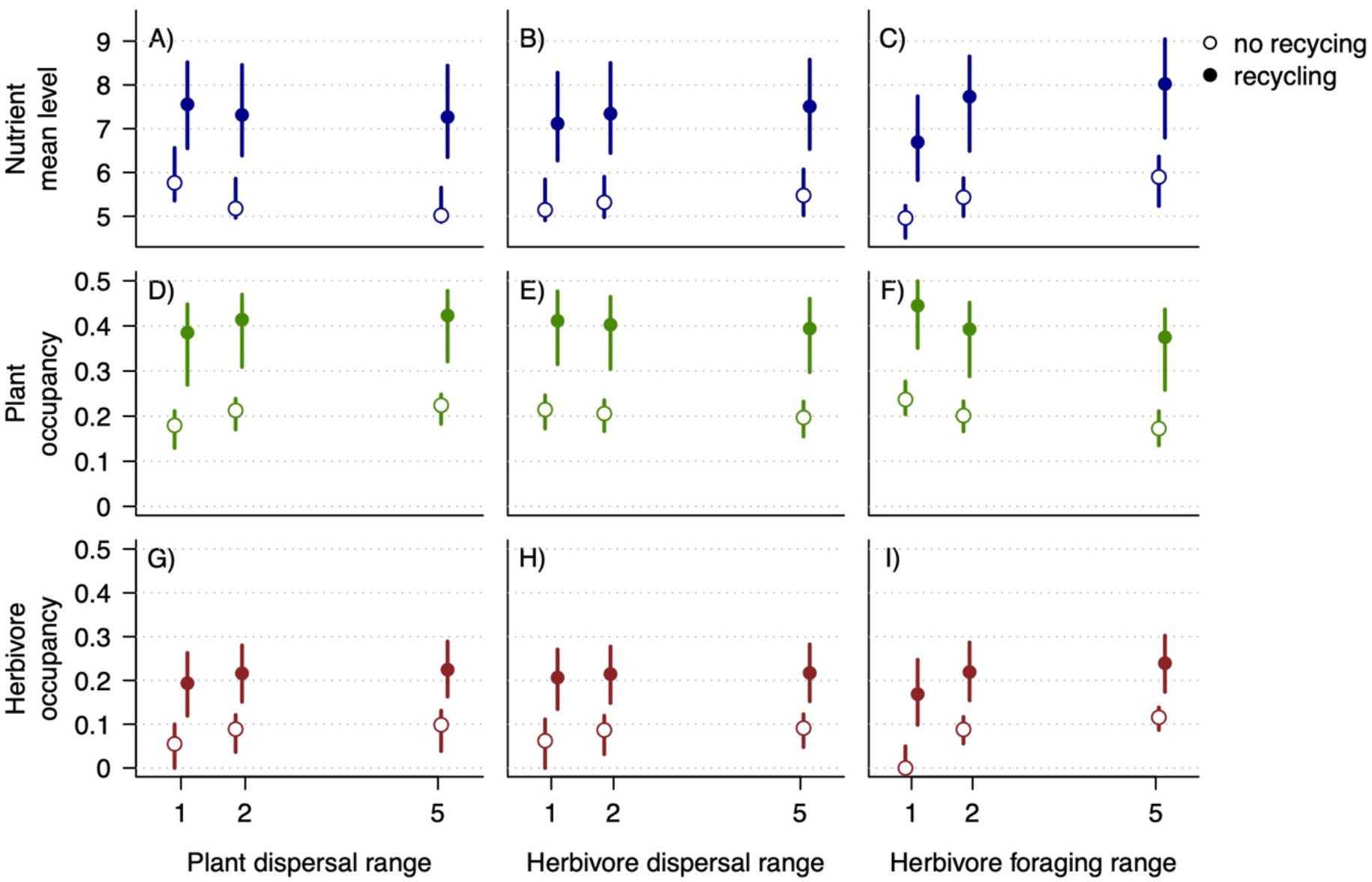
Mean Nutrient value (A-C) and Plant (D-F) and Herbivore (G-I) occupancies for varying plant dispersal, *Π*_*P*_, herbivore dispersal, *Π*_*H*_, and herbivore foraging, π_*H*_, ranges. Open and solid circles represent simulations without or with recycling (*σ*_*P*_ = *σ*_*H*_ = 0 or *σ*_*P*_ = *σ*_*H*_ ∈ {1,2}, respectively). Blue, green and red colors correspond to nutrient, plant, and herbivore, respectively. Points and bars give the quartiles. All colonization-extinction scenarios are pooled. *θ* = 0.5. See Table 3 for other parameter values.

### 4.2. Plant dispersal is a major driver of spatial structure across trophic levels

Plant dispersal range is the dominant driver of spatial clustering of the plants themselves, but also of nutrients, and it feeds back to herbivores (Figs 3 and 4). Herbivore dispersal range only affects its own spatial clustering, and foraging range has little effect on spatial structures (Fig. 3G-I).

**Figure 3.**
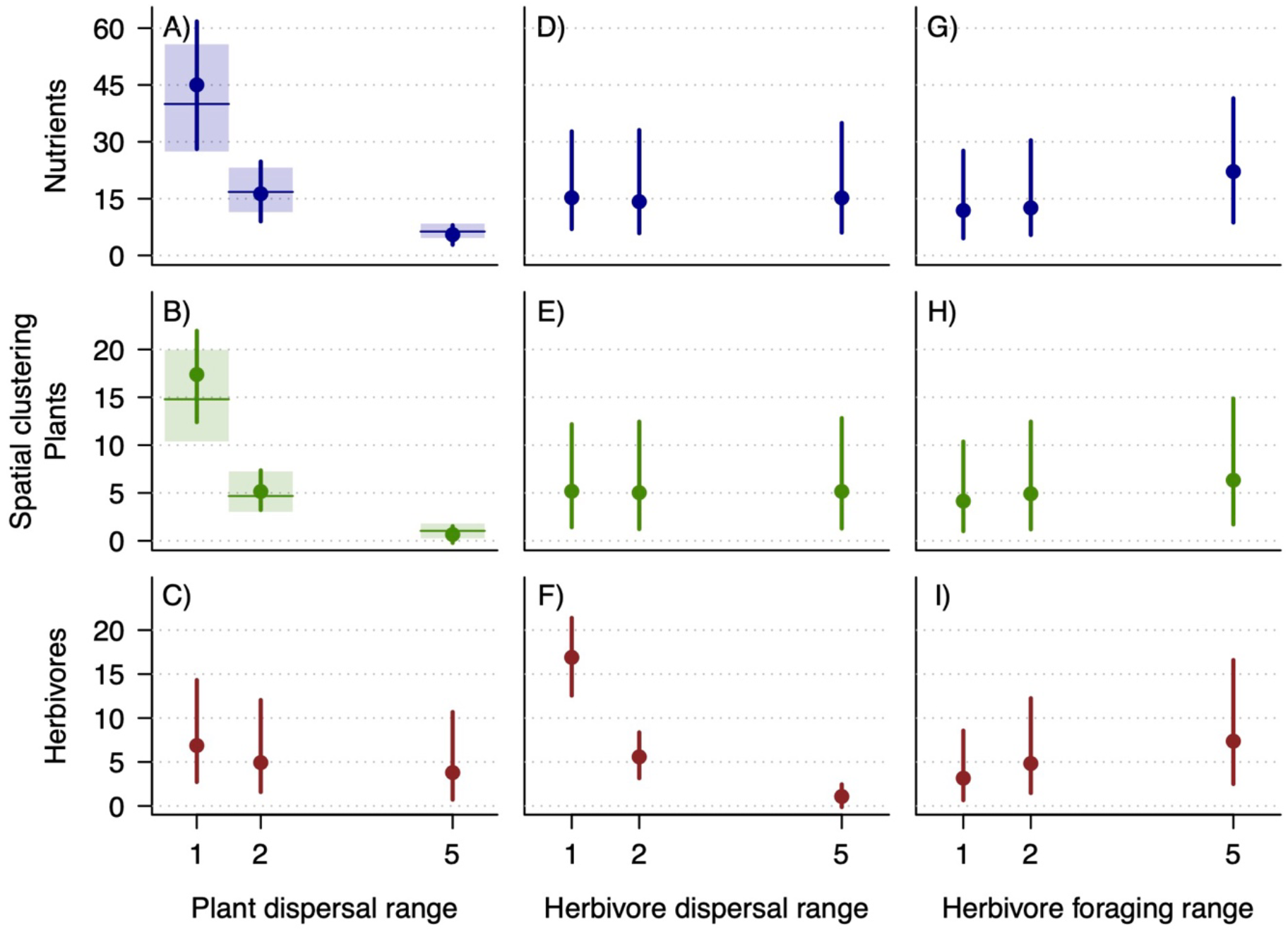
Spatial clustering of nutrients, and plant and herbivore populations for varying plant dispersal (A-C), *Π*_*P*_, herbivore dispersal (D-F), *Π*_*H*_, and herbivore foraging (G-I), *π*_*H*_, ranges. Spatial clustering is measured as the Z-score of the Moran index (spatial autocorrelation). Moran indices quantify near-neighbor correlation and increases as neighboring cells are more similar and the z-score allows to control for variations in occupancy among scenarios. Points and bars give the quartiles for scenarios with herbivores. Horizontal lines and shadow areas give the quartiles for scenarios without herbivores. All recycling and colonization-extinction scenarios are pooled. *θ* = 0.5. See Table 3 for other parameter values.

**Figure 4.**
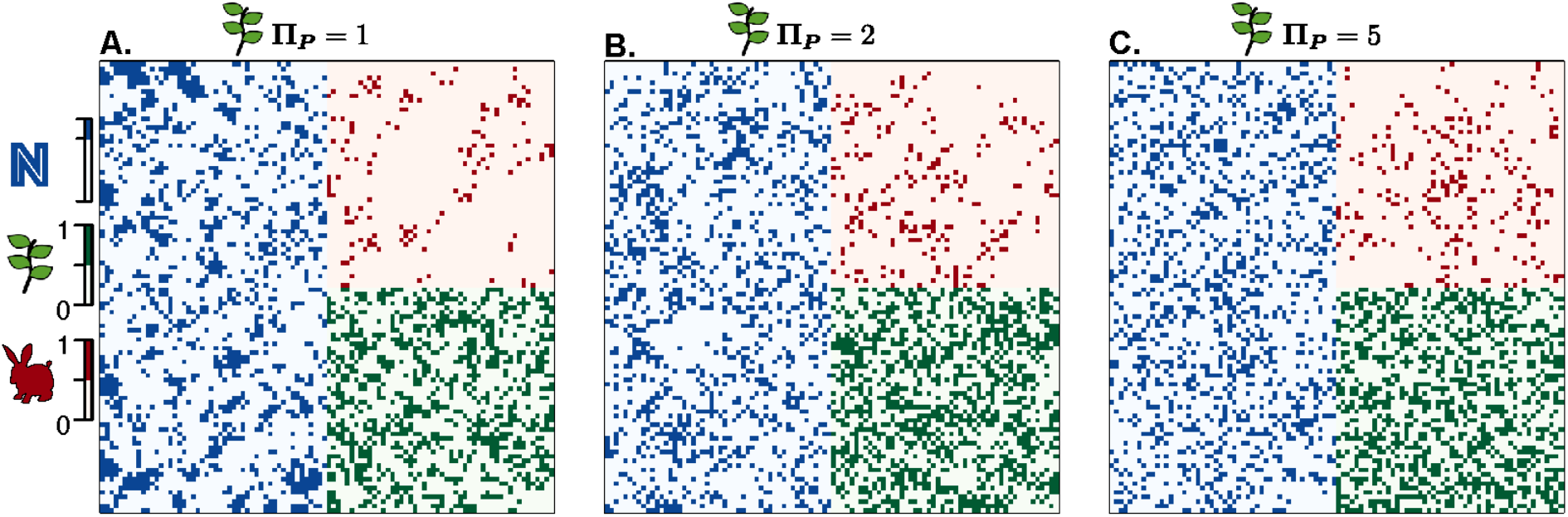
**Nutrient availability and plant and herbivore occupancy for 3 landscapes with varying plant dispersal ranges** that are representative of the mean Moran N z-scores of left panels in figure 3. Panels show distributions of nutrients which level is in the highest quartile in the left ½ of the landscapes (blue pixels; see Fig. S3 for continuous values), plant occupancy in the lower right ¼ (green pixels), and herbivore occupancy in the upper right ¼ (red pixels) of the landscape. Parameter values *σ*_*P*_ = *σ*_*H*_ = 1, *ρ*_*P*_ = 1.6, *e*_*P*0_ = 0.4, *ρ*_*H*_ = 6, *e*_*H*0_ = 0.4, *Π*_*H*_ = 2, π_*H*_ = 1. *θ* = 0.5. See Table 3 for other parameter values.

As plant dispersal range increases, the spatial clustering of nutrients, plant, and herbivore populations decreases (Fig. 3A). Local dispersal (small *Π*_*p*_) creates large clusters of plant populations (e.g., Fig. 4A) that largely deplete local resources, leading to nutrients being aggregated in unoccupied cells. The spatial aggregation of plants also leads to a certain level of aggregation in herbivore populations, though the aggregation is weaker than at the plant level, as herbivores that have large foraging ranges are less affected by plant clustering. The clustering of plants and nutrients can be observed in the absence of herbivores as well (shaded areas in Fig. 3A-B). Overall, we observe a strong effect of plant dispersal range in all simulation results (i.e., across different scenarios). This highlights the importance of plant dispersal in spatially structuring nutrients that might mask a potential effect of herbivores, which we then expect to be detectable only within a given plant dispersal range.

### 4.3. Conditions for herbivores to affect nutrient spatial structure

Herbivores need recycling to affect nutrient patterns. In our simulations without recycling, nutrient levels as well as plant and herbivore occupancies are globally low, but all three increase through the fertilizing effect of recycling (Fig. 2). As a corollary, the spatial clustering of nutrients (and of both species) decreases globally with recycling (Fig. 5, S4). This effect occurs because recycling redistributes nutrients in the landscape and thereby fills nutrient cold spots that had emerged due to depletion by plants. This also implies that herbivore occupancy becomes higher, so if there is a numerical effect of herbivores on plant (i.e., top-down control) or on nutrient spatial structure, we expect it to be more visible in the presence of recycling. Consistent with this expectation, we observe a stronger positive effect of the herbivore foraging range on the spatial structuring of resources (and plants) with recycling than without, at a given plant dispersal range (Fig. 5A-C). However, herbivore dispersal range has no effect on nutrient (and plant) clustering, whatever the recycling level, although its variation generates a strong gradient of spatial clustering in herbivore populations (Fig. 5D-F). This counter intuitive result shows that the major path by which herbivore populations affect the distribution of nutrients in our model is not the direct imprint of their own spatial distribution via recycling but rather the indirect effect through limiting plant colonization within their foraging range.

**Figure 5.**
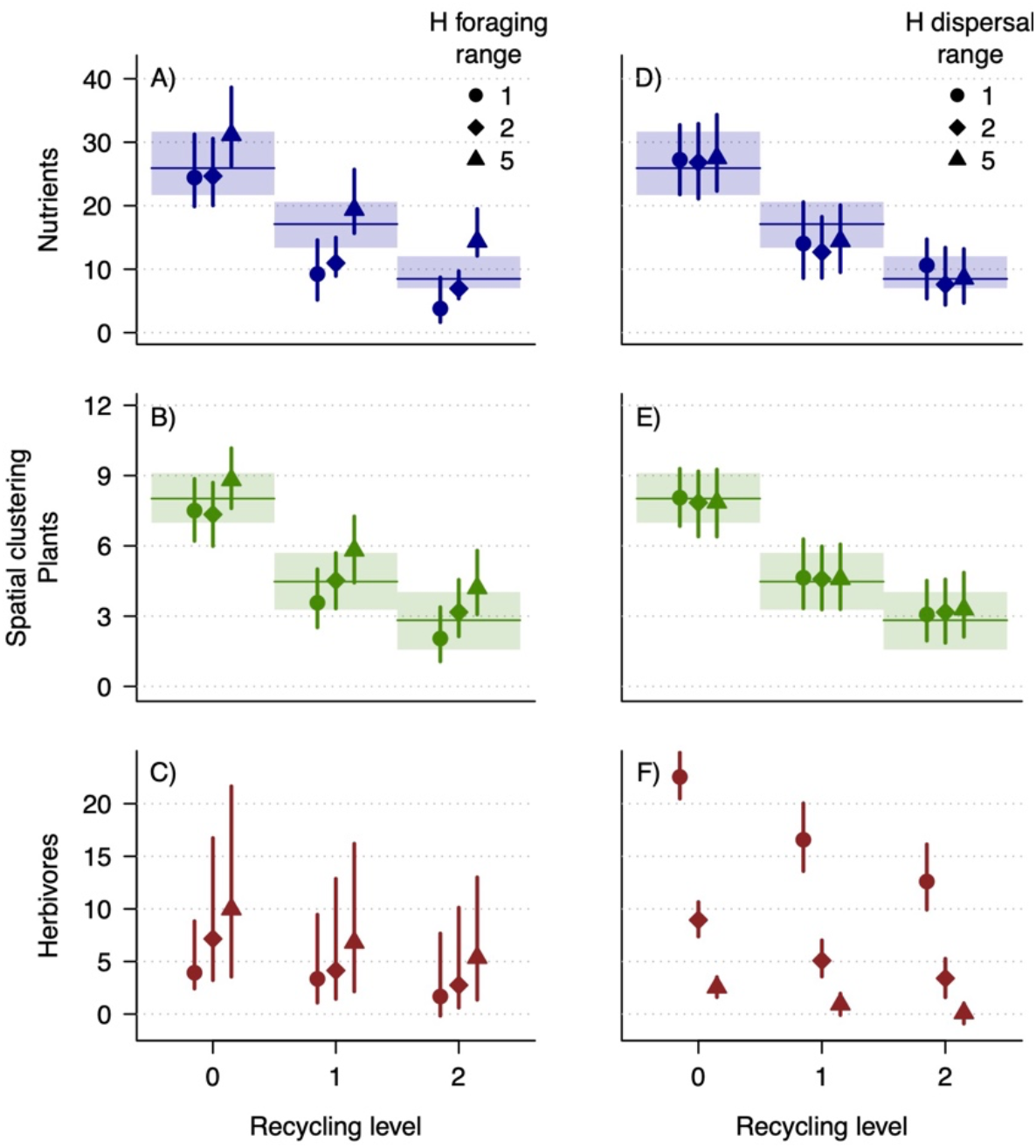
Effect of herbivore foraging, *π*_*H*_, (A-C) and dispersal, *Π*_*H*_, (D-F) ranges (shapes) on the spatial clustering of nutrients, Plant and Herbivore populations for varying recycling levels (x-axes) at a given dispersal range of the plant populations: *Π*_*P*_ = 2. Spatial clustering is measured as the Z-scored of the Moran index (spatial autocorrelation). Points and bars give the quartiles. Horizontal lines and shadow areas give the quartiles for scenarios without herbivores. All colonization-extinction scenarios are pooled. *θ* = 0.5. See Table 3 for other parameter values.

### 4.4. Foraging range dynamically and indirectly affects spatial structure

So, what is driving the increase in spatial clustering of all trophic levels with increasing herbivore foraging range (Fig. 5A-C)? This effect is mediated by herbivore population persistence and dispersal as follows: when the foraging range of herbivores is small, there is a small probability of herbivores having plants in their foraging range, and the population quickly goes extinct (Fig. 6A). Fast extinction implies reduced opportunities for dispersal, and hence lower occupancy and lower top-down control of plants. By contrast, when the foraging range of herbivores is larger, there is a higher probability of herbivores having plants in their range. Regardless of possible plant extinctions, higher foraging range provides herbivores a more reliable resource supply and their survival is prolonged (Fig. 6A).

**Figure 6.**
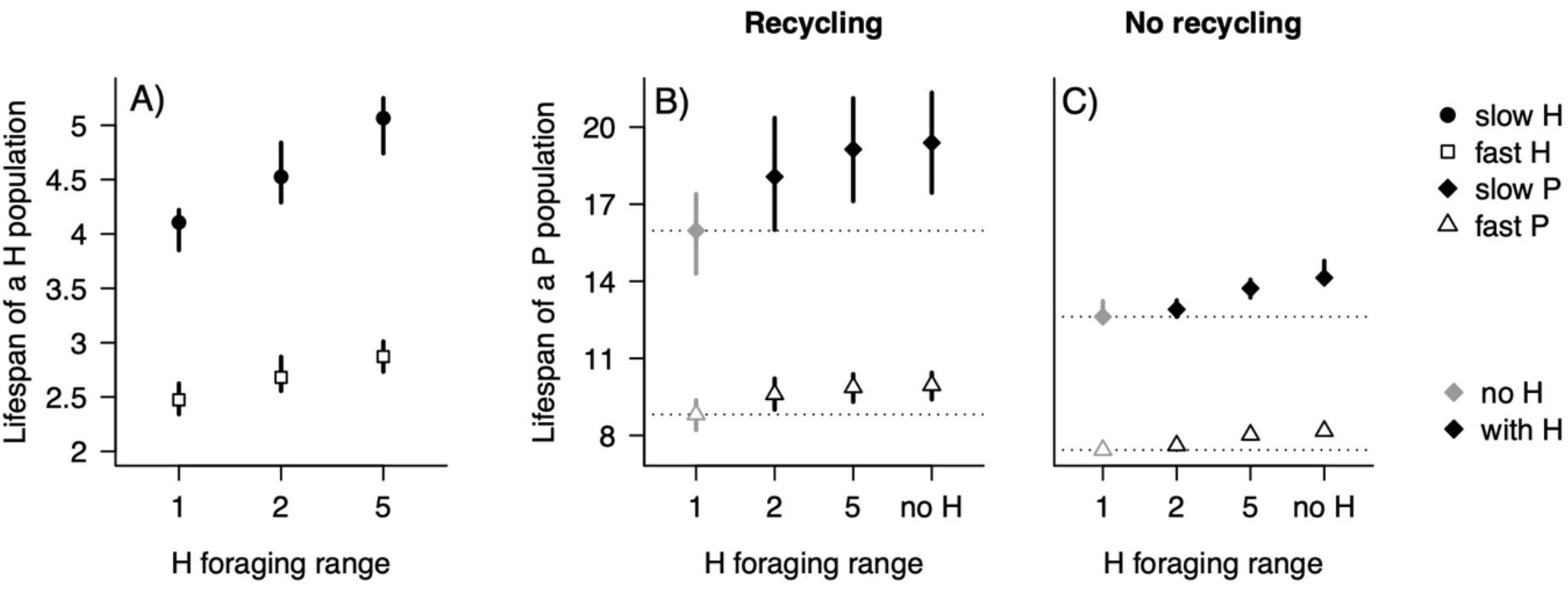
Number of timesteps a population stays in a given cell before extinction for varying herbivore foraging range, *π*_*H*_. Points and bars give the quartiles. A) Herbivore populations, B-C) Plant populations for the cases B) with or C) without recycling (*σ*_*P*_ = *σ*_*H*_ = 0 or *σ*_*P*_ = *σ*_*H*_ ∈ {1,2}, respectively). We distinguish populations which have slow (solid shapes) and fast (open shapes) colonization-extinction dynamics as it obviously drives population lifespan (see Table 4 for parameter values). Grey symbols and dotted lines in B) and C) show the lifespan of a plant population for corresponding scenarios without herbivores. Scenarios of all dispersal ranges (*Π*_*P*_, *Π*_*H*_) are pooled.

Plant populations also have longer lifespans in the presence of herbivores than in their absence, and with increasing herbivore foraging range (Fig. 6B). As the effect persists in absence of recycling (Fig. 6C), it can only result from herbivory pressure relaxing the exploitative competition among plant populations by decreasing plant occupancy and thus increasing N. Longer lifespans imply more time to create hot and cold spots locally via depletion and recycling, which contributes to generating nutrient heterogeneity when associated with aggregation mechanisms. Aggregation could come from herbivore local dispersal (small *Π*_*H*_) as daughter populations will cluster. In turn, these local persistent herbivore clusters should impact nutrients, which we indeed observe (Fig. 7A-C). The clustering of nutrient is, however, even stronger when herbivores disperse globally while no spatial structure occurs at their level (Fig. 7D-F). The only possible explanation is that the homogenous distribution of herbivore populations under global dispersal create patches under strong herbivory pressure where plants cannot colonize when the foraging range is large (Fig. 7G).

**Figure 7.**
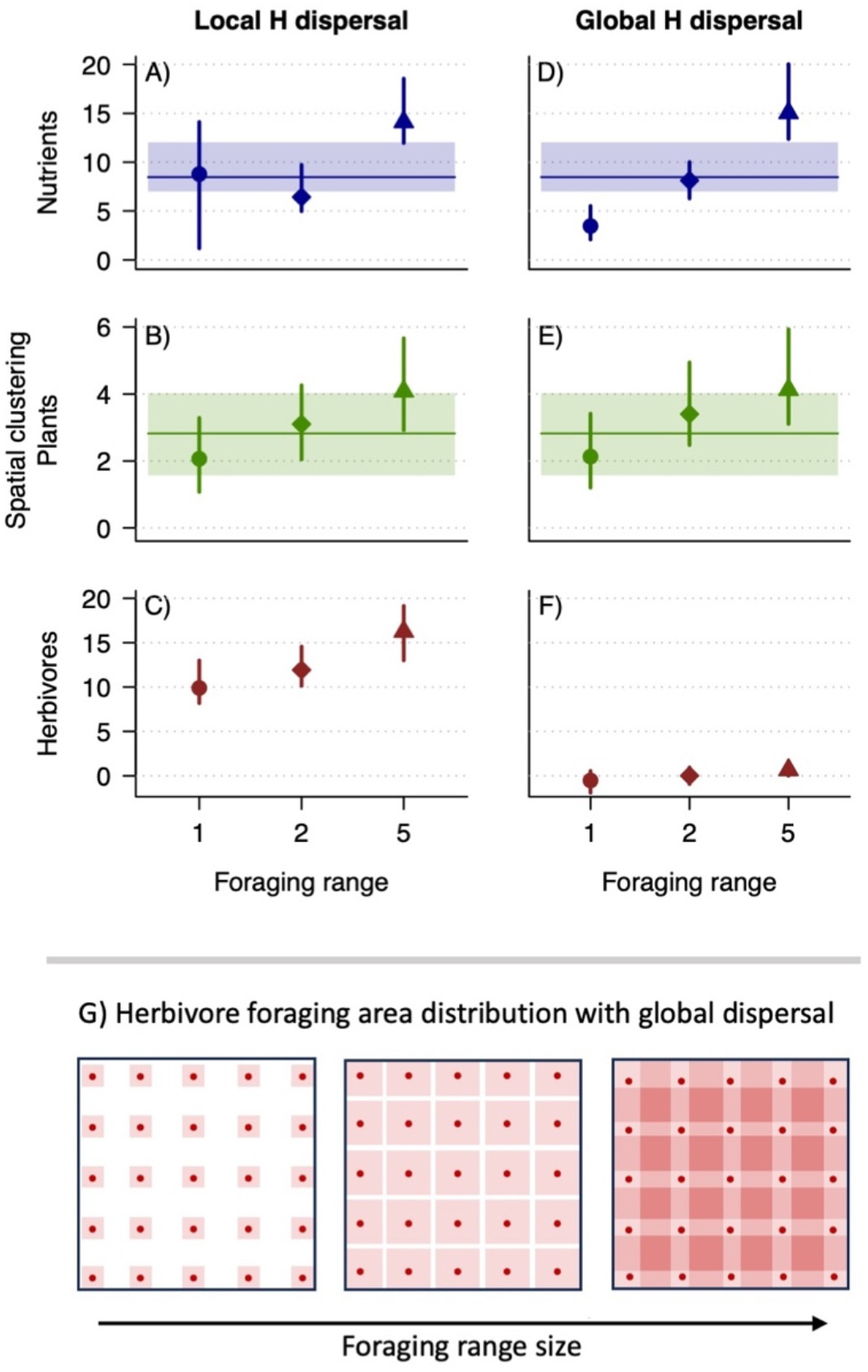
Effect of Herbivore foraging ranges *π*_*H*_, at local (A-C, *Π*_*H*_ = 1) versus global herbivore dispersal range (D-F, *Π*_*H*_ = 5) on the spatial clustering of Nutrients, Plant and Herbivore populations at plant dispersal range *Π*_*P*_ = 2 and high recycling level (*σ*_*P*_ = *σ*_*H*_ = 2). Spatial clustering is measured as the Z-scored of the Moran index (spatial autocorrelation). Points and bars give the quartiles. Horizontal lines and shadow areas give the quartiles for scenarios without herbivores. All plant colonization-extinction scenarios are pooled. Panel (G) illustrates the distribution of herbivore foraging areas (squares) and their potential overlap with increasing range size when populations (points) are equally spaced (global dispersal). The darker the color, the higher the herbivory pressure, the lower the probability of plant colonization.

## 5. Discussion

We derived a mechanistic model to study how feedbacks in a nutrient-plant-herbivore meta-ecosystem affect spatial patterns of nutrients. We demonstrate how spatial heterogeneity in nutrients at the landscape scale can emerge solely from organisms*’* consumption, movement, and recycling processes, without any underlying environmental constraint. We find dispersal range of plants to be a major determinant of nutrient spatial heterogeneity by driving the spatial distribution of plants and consequentially nutrient uptake. Effects of herbivores on nutrient distribution are mainly mediated by how herbivore consumption constrains the spatial distribution of plants rather than by a direct imprint of herbivore distribution via nutrient recycling. However, recycling plays an important role in the capacity of herbivores to significantly affect nutrient distribution, by making the system productive enough to sustain high herbivore occupancy. Herbivore-induced nutrient patchiness then emerges from a combination of temporal and spatial mechanisms appearing with large herbivore foraging ranges. Large herbivore foraging ranges both allow plant and herbivore populations to occupy the same place, and thus to affect local resources a longer time, and create areas of high herbivory pressure unfavorable to plant colonization where nutrients can accumulate. We discuss below these findings and their relevance for real ecosystems in the light of current empirical literature.

### Plant spatial patterns and nutrient distribution

In our spatially-explicit occupancy model, spatial heterogeneity in nutrients mainly emerges from plant dispersal limitation. When plant populations disperse locally, vegetation aggregates in patches and nutrients are locally depleted. Plants thus induce landscape-scale patchiness in nutrients reflecting their own patchy spatial distribution. The ability of individual plants to generate fine-scale heterogeneity in soil has long been recognized (Waring et al., 2015). At the landscape scale, alpine and arid ecosystems provide iconic examples of vegetation patterns reflected in soil properties through physical feedbacks between plant and soil processes (Klausmeier, 1999). While our model does not integrate these physical feedbacks, instead, it demonstrates that nutrient heterogeneity can emerge at the landscape scale solely from plant colonization-extinction dynamics and local nutrient uptake and recycling. Analogous empirical examples include disturbed forests where tree mortality creates gaps that are temporarily richer in nutrients than surrounding forested patches due to uptake release, before recolonization occurs (Tong et al., 2024). Beyond sites with contrasting open and vegetated areas, nutrient heterogeneity may also emerge within diverse vegetation, when plant species with a small dispersal range, which makes them clump, have contrasting effects on soil nutrients. For instance, in a calibrated tree succession model of boreal forests, local dispersal of spruce, which has a low-quality litter compared to early-succession trees, produced soil nitrogen patchiness in an otherwise homogenous environment (Pastor et al., 1999); and, as in our model, this patchiness is dampened when seeds disperse globally. In the same line, Wilcots et al. (2019) found that rhizobia nitrogen-fixing plants have, on average, significantly larger and more biotically-dispersed seeds than non-fixing plants, which predispose them to be spatially clumped, with the potential of forming nutrient hotspots due to their nitrogen-rich leaf litter. Nutrient spatial patterns can also be induced by other sessile organisms than primary producers, such as filter feeders, with mussels being able to aggregate in large beds and creating biogeochemical hotspots due to their important impact on nutrient recycling (Atkinson & Vaughn, 2015). Overall, the ingredients making some basal species a source of spatial clusters in nutrients is a combination of aggregation mechanisms and contrasting effects on nutrient fluxes compared to other present basal species.

### Herbivores influence nutrient patchiness through their effect on plant distribution

On top of this central role of primary producers, our model first stresses the importance of ecosystem productivity for herbivores to be abundant enough to have any effect –positive or negative– on nutrient patchiness. This should be especially important in terrestrial systems, where low-quality food maintain herbivores at low biomass compared to plants (Bar-On et al., 2018). Second, in our model, the effects of herbivores on nutrient spatial structure are mainly mediated by the spatial constraints exerted on the spatial distribution of plants rather than by herbivore recycling. A similar finding was found, for instance, by Augustine and Frank (2001) with large herbivores in Yellowstone Park affecting soil nitrogen heterogeneity mainly through their top-down effect on plants despite the important N return to soil in dung and urine. Actually, in this study, herbivores decreased nitrogen spatial heterogeneity at small spatial scale (0.1-1m) compared to exclosures by promoting fine-scale plant diversity, and increased it at large scale (30m) due to grazing preferences linked to field slopes. In our model, nutrient hotspots are created without topographic variability, in areas relaxed from plant depletion due to high herbivory pressure. A global meta-analysis shows that mega-herbivores indeed tend to promote open and semi-open habitats by increasing bare ground and soil compaction and decreasing plant biomass at the plot scale, while average effects on soil nutrient varies. At a higher –inter-plot– scale, only the largest herbivores (> 100 kg) promote diversity in vegetation structure and nitrogen heterogeneity (Trepel et al., 2024). This aligns with our result showing that nutrient heterogeneity tends to increase only for large foraging ranges, which generally correspond to larger animals (Straus et al., 2024). Further theoretical investigation of the links between herbivore body size, movement scales and recycling effects should refine our understanding of the scale of the feedback between herbivores and nutrient spatial heterogeneity, and the underlying mechanisms (Ferraro & Lienau, 2025).

### Dynamical landscapes and the persistence of nutrient patches

In our model, the spatial distribution of populations is dynamic and varies in time along with species colonization-extinction processes, and consequently nutrient patches. One surprising result of our model is that aggregations of herbivore populations does not guarantee the emergence of nutrient clusters because herbivore aggregations may be too ephemeral to have any impact on nutrient distribution. On the contrary, herbivores*’* maximum impact on nutrient distribution occurred when they disperse globally –and thus do not form clusters– which allow them to better track plant hot spots and reach higher occupancies. This, in combination with large foraging ranges, increases the persistence of their populations, as well as those of plants by reducing intra-plant competition. Higher abundances and longer population lifespans are thus key ingredients for herbivores to locally affect nutrient distribution. The time populations persist in one place may depend on various mechanisms, beyond the herbivory pressure and external perturbations considered here, such as species life-history traits modulating individual*’*s lifespan (e.g., grasses versus trees). Interestingly, we note that the simple processes accounted for in our model –i.e., consumption, recycling, extinction and colonization– are not sufficient to create perennial nutrient hot spots when the background nutrient level is spatially homogenous. The empirical literature tells us that in absence of some topographical heterogeneity which traps nutrients, creating permanent nutrient hot spots needs some spatial heterogeneity in the biological processes of consumption or recycling, or some self-reinforcement processes whereby species modify the physical environment in a way that concentrates locally nutrients, such as with beavers or mussel beds. Non-spatially homogenous consumption can result, for instance, from social behavior (Ferraro et al., 2022), selective plant foraging by herbivores which can create patches of plants having different impacts on soil nutrients (Adler et al., 2001; Bloor & Pottier, 2014), or presence of predators constraining the use of habitat by herbivores (Teckentrup et al., 2018), for instance depending on their size, with some consequences on nutrient distribution. Indeed, Le Roux et al. (2020) showed that in African savannas, smaller herbivores graze more in open areas in the presence of predators than the largest herbivores, as they need to see predators approaching. Since smaller herbivores also recycle more phosphorus than larger ones, the latter retaining more phosphorus in their larger P-rich skeleton, then open areas receive proportionally more fecal phosphorus from herbivores than forested areas in the presence of predators. Another common mechanism leading to perennial nutrient clusters involves spatially localized recycling, where animals defecate, urinate or deposit natal fluids in specific places to avoid parasites or predators when foraging or resting in the same place. This behavior is observed for instance in monkeys (Feeley, 2005), cattle grazing in alpine pastures (Jewell et al., 2007), or caribou calving areas (Ferraro et al., 2024). Models integrating such heterogeneity in spatial behavior and biological processes would allow to extend our understanding of biotic-abiotic feedbacks leading to the emergence of biologically-induced perennial nutrient heterogeneity.

## Conclusions

Overall, our model identifies minimal-ingredient mechanisms creating biologically-induced nutrient patchiness at the landscape scale. In absence of any spatial heterogeneity imposed by the environment or by organism behaviors, nutrient hot spots could emerge through the interaction of animal and plant consumption, recycling, and movement. In these minimal conditions, although plant dispersal should be the dominant driver of nutrient patchiness, we demonstrate herbivores can also have an impact in sufficiently productive systems, when their large dispersal and foraging ranges make them efficient to track plant hot spots and prolonged the local persistence of their populations. By showing the conditions for animals to drive nutrient heterogeneity, our results add new theoretical predictions to the emerging field of zoogeochemistry (Leroux & Schmitz, 2025).

Understanding the dynamical biotic-abiotic feedback, and the scales at which it happens, is an essential condition to building a more mechanistic view of how both abiotic and biotic landscapes are altered by humans. Conversely, human activities, especially land-use changes and species introductions, also lead to large changes in the distribution and space use of many species (Riotte-Lambert & Matthiopoulos, 2020; Tucker et al., 2018) and to large decreases in their population sizes (Dirzo et al., 2014). In this context, understanding the dynamic biotic-abiotic feedback loop could allow us to better forecast future resource distributions and their implications for ecosystem functions and services.

## Acknowledgements

This research is product of the RED-BIO group funded by the synthesis center CESAB of the French Foundation for Research on Biodiversity (FRB; www.fondationbiodiversite.fr) and the CIEE (https://www.ciee-icee.ca/).

## Author contributions

Isabelle Gounand, Nicolas Loeuille, Emanuel Fronhofer, Eric Harvey, Sonia Kéfi, Shawn Leroux, Chelsea Little, and François Massol conceived the study and developed the model. Isabelle Gounand coded the model and run the simulations with the help of Camille Saade. All authors analyzed the outputs and contributed critically to the writing of the manuscript.

## Data availability

the julia source code generating the data, and the R scripts generating the figures will be made available on zotero XX.

