## Supplementary Material for "Nutrient heterogeneity emerges from dynamical abiotic-biotic feedback in a spatially explicit plant-herbivore occupancy model"

### Appendix 1: Supplementary methods

#### 1.1 General overview

We describe a landscape with a spatially explicit occupancy model including an abiotic resource (e.g., nitrogen or phosphorus), hereafter nutrient $N$ , populations of primary producers and consumers, thereafter respectively referred to as resource, plants $P$ and herbivores $H$. We do not model local plant and herbivore demography (i.e., mass transfers) but rather the effect of plant (or herbivore) presence on herbivore (or plant) colonization-extinction dynamics. The landscape consists of a square lattice of 100 x 100 cells, where each grid cell contains a quantitative level of resource, and might, or might not, be occupied by a population of plants, a population of herbivores, or both (Fig. 1). As such, each cell or any combinations of cells capture interactions between abiotic and biotic components and are therefore, ecosystems.

In a given cell, the resource exchanges matter with a background resource pool (e.g., atmospheric deposition, rock weathering, leaching), and diffuses into its eight neighboring cells (Fig. 1a). One cell corresponds to the foraging range of a plant population. A plant population depletes the resource locally (consumption), and may also replenish the resource locally through local recycling of detritus into the local resource pool (catabolism and mortality). Plant populations may colonize cells without plants within their dispersal range, (see below) or go extinct. Resource amount affects plant colonization and extinction dynamics by increasing the chance of a cell being colonized, and decreasing the chance of extinction.

The number of herbivore populations around a given cell also decreases the chance of plant colonization due to their foraging activity, i.e. they only affect plant populations in their foraging range (Fig. 1b). Herbivore populations feed upon the plant populations present in their foraging range. Herbivore populations can colonize empty cells within their dispersal range, with a higher probability if the new foraging range contains more plant populations and fewer herbivore ones. Herbivore populations can go randomly extinct (e.g. due to catastrophic events such as floods or disease outbreaks), but with a higher probability when their foraging area becomes poor in plant populations (due to food shortage). Recycling of herbivore biomass follows comparable rules as for plants, but detritus is spread equally across the whole foraging area. Note that local colonization/extinction probabilities of herbivores depend not on the presence of plants in the local cell, but on the total plant occupancies within its foraging range (Fig.1c).

Plants and herbivores are thus characterized, among other features, by the respective sizes of their foraging ($\pi_P$, $\pi_H$) and dispersal ($\Pi_P$ and $\Pi_H$) ranges, with $\pi_P$ set to $0$ corresponding to the plant foraging in one cell only ($\pi$ parameters are defined so that foraging areas are squares centered on their focal cell, with sides equal to $2\pi + 1$; in the same way, the area in which populations can disperse is defined as a square of side $2\Pi + 1$). We aim at understanding how these range sizes affect the spatial distribution of abiotic resources.

#### 1.2 Resource dynamics

We broadly follow classic cellular automata models for the resource layer with deterministic dynamics in discrete time (Dieckmann et al. 2000, chap. 6). In the absence of plants, the resource value in a given cell is defined by exchanges with a background resource pool (chemostat processes), and with the surrounding cells (diffusion), which tends to homogenize resource levels in the landscape and to equalize it with the background level. Plant populations decrease the resource in the local cell while any local recycling from plant or herbivore populations increases the value of the local resource. Please note that, since the model focuses on populations, not individuals, recycling occurs through both the detritus produced by excretion, dead leaves, etc. and those associated with dead individual plants or herbivores.

More specifically, each cell is characterized by a resource value, $N_i$ which changes at each timestep $t$ ($\Delta N_i(t) = \tau\phi_i(t), \tau$ being the size of a timestep) with a rate $\phi_i(t)$ equal to the sum of effects due to the five processes listed above (diffusion, chemostat, plant uptake, plant recycling, and herbivore recycling; see Fig. 1a)

$$\frac{\Delta N_i(t)}{\tau} = \phi_i(t) = \phi_i^{\delta}(t) + \phi_i^{\xi}(t) + \phi_i^{\upsilon}(t) + \phi_i^{\sigma_P}(t) + \phi_i^{\sigma_H}(t)$$

where:

- $\phi_i^{\delta}(t) = -\delta(N_i(t) - N_0)$ is the contribution of the chemostat process between the background source of nutrient pools (at value $N_0$) and the focal cell, which tends to come back to the background level, arbitrarily set to 10 as a reference value, with a strength modulated by rate $\delta$;

- $\phi_i^{\xi}(t) = -\xi\left(N_i(t) - \frac{1}{|V_i|}\sum_{k\in V_i} N_k(t)\right)$ is the contribution of the diffusion process between cell $i$ $i$ and the eight cells belonging to the resource-neighborhood $V_i$ of cell $i$; this effect tends to spatially homogenize resource levels and it increases with the difference between $N_i$ and the mean level of the eight nearest neighbors, modulated by the diffusion rate $\xi$ ;

- $\phi_i^{\upsilon}(t) = -\upsilon P_i(t) max[N_i(t) - N^*, 0]$ is the contribution of the plant uptake process, which tends to have $N_i$ decrease towards lowest resource level $N^*$, set to 0, when a plant population occupies the cell ($P_i = 1$), with a strength depending on the depletion rate $\upsilon$ and the distance $N_i$ is from $N^*$. The maximum function ensures that $N_i$ does not go below $N^*$;

- $\phi_i^{\sigma_P}(t) = \sigma_P P_i(t-1)$ is the contribution of plant populations to recycling in cell $i$ at time t, when a plant was present at time $t-1$ ($P_i(t-1)$ equals 1 when a plant population is present and 0 when absent), and is modulated by the plant recycling rate $\sigma_P$;

- $\phi_i^{\sigma_H}(t) = \sigma_H \eta_i(t-1)$ is the contribution of herbivore populations to recycling in cell $i$ at time $t$, when at least one herbivore population was present at time $t-1$ in the foraging range around the focal cell $i$, and is modulated by the herbivore recycling rate $\sigma_H$, with experienced herbivore recycling computed as the number of cells occupied by an herbivore population in the foraging range around cell $i$, $Z_i$, and cell $i$ itself at time $t-1$, divided by the number of cells in the foraging range: $\eta_i(t) = \frac{1}{1+|Z_i|} \sum_{k \in Z_i \cup \{i\}} H_k(t)$ , as we assume detritus from herbivore populations to be spread equally over the whole foraging range. The set $Z_i$ corresponds to all cells from which herbivore populations could feed upon cell $i$ as part of their foraging range.

#### 1. 3 Colonization-extinction dynamics

Each cell can harbor one population of plants, one population of herbivores, or both. A plant population can experience herbivory from all herbivore populations whose foraging range covers the plant's cell. Plant and herbivore occupancies are governed by stochastic processes of extinction and colonization, which depend notably on food availability, as well as on herbivory pressure for plant colonization. Based on their rates, a number of events can take place, which fall into four categories:

- *Plant population extinction*: an existing plant population (i.e. a cell $i$ in which $P_i = 1$) dies out i..e. $P_i$ goes from 1 (occupied) to 0 (empty cell). Each existing plant population has a rate $e_{Pi}$ of going extinct which combines a basal rate due to possible perturbations $e_{P0}$, with a dependency on the resource level in the cell, $N_i$, according to an inverse exponential function: $e_{Pi} = e_{P0} + \epsilon e^{-\beta_P(R_i - R^*)}$; the extinction rate increases exponentially when , the parameter $\beta_P$ regulating the steepness of the exponential.

- *Herbivore population extinction*: an existing herbivore population dies out ($H_i$ goes from 1 to 0) at a rate $e_H$, which similarly combines a basal rate due to possible perturbations $e_{H0}$ with a dependency on food availability in the foraging area around the cell, $\lambda_i$, formalized by an inverse exponential function (Fig. 1c). Extinction rate becomes maximal when $\lambda_i$ approaches $\lambda^*$, the minimal food requirement for herbivores. $\lambda_i$ includes competition with other herbivores present in the foraging range of the population $Z_i$. It, is calculated as the sum of plant populations in the foraging range divided by the number of herbivore populations able to feed on it (Fig. 1c):

- *Plant population colonization*: a cell devoid of plants can be colonized by other plant populations within its dispersal range ($P_i$ goes from 0 to 1). The colonization rate into patch *i*, $c_{P,i}$, increases with the abundance of potential colonizer populations and, also with resource availability, but decreases with herbivore pressure, according to the following expression: $c_{P,i} = c_{P0} \times f_P(R_i) \times (1 - a\kappa_i^{\theta})$(Fig. 1b), where $c_{P0}$ is the basal colonization rate of plant populations, which depends on both plant colonization effort, $\rho_P$, and plant occupancy in the plant dispersal area around cell *i*, $U_i$: $c_{P0} = \frac{\sum_{k \in U_i} P_k \rho_P}{|U_i|}$; function $f_P(N_i)$ gives the resource dependency of the probability that the cell can be colonized, which increases toward 1 according to a logit function (see Fig. 1b) showing a logistic shape, with steepness being regulated by $\alpha_P$ and inflexion point by $N_{col}$; $\kappa_i$ is a measure of herbivore pressure in site $i$ given by the relative abundance of herbivore populations inhabiting cells within the foraging neighborhood $Z_i$ of cell $i$, expressed as: $\kappa_i = \frac{1}{1 + |Z_i|} \sum_{k \in Z_i \cup \{i\}} H_k(t)$. Coefficients $a$ and $\theta$ are regulating the strength and non-linearity, respectively, of the negative effect of herbivore pressure on plant colonization rate (here chosen to be linear, i.e., Fig. 1b).

- *Herbivore population colonization*: a cell devoid of herbivores can be colonized by herbivore populations ($H_i$ goes from 0 to 1) within its dispersal range. Colonization probability increases with the number of potential disperser populations and with food availability: $c_{H,i} = c_{H0} \times f_H(\lambda_i)$. The basal colonization rate, $c_{H0}$, depends on both herbivore colonization effort, $\rho_H$, and herbivore occupancy in the corresponding dispersal area $W_i$: $c_{H0} = \frac{\sum_{k \in W_i} H_k \rho_h}{|W_i|}$; the function $f_H(\lambda_i)$ gives the probability that the site can be colonized based on the available food in the foraging range around cell $i$; the function increases toward 1 according to a logit function showing a logistic shape (Fig. 1c), with steepness being regulated by $\alpha_H$ and inflexion point by $\lambda_{col}$.

#### 1. 4 Simulation dynamics

We assume colonization and extinction events to follow Poisson processes (i.e., Markovian processes in continuous time occurring with given rates). Instead of resorting to the classic Gillespie algorithm to simulate these processes, we used the tau-leaping algorithm proposed by Gillespie (2001) with a constant time step (tau-leap) to easily coordinate discrete biotic events with discrete-time resource dynamics. The idea of the tau-leaping method is to approximate Gillespie's algorithm on regular intervals $\tau$ that are sufficiently short so that the propensities of the different event types (rates summed over the grid) do not change much during the interval. During a tau-leap ($\tau$), a number of random events can take place, which fall into the four categories described above (colonization or extinction of a plant or an herbivore population). These events are drawn using the following procedure:

- At each time step, we calculate the rates of each event in each cell of the landscape, as well as the propensity of each type of event by summing the rates at which each type of event can occur in all sites (e.g., propensity of herbivore extinctions is $\sum e_{H,i}$, with $i \in lattice$).
- For each type of event, the number of events happening is drawn from a Poisson distribution with a mean equal to the propensity of this type of events multiplied by the time step size $\tau$.
- The cells in which events happen are then drawn from a multinomial distribution, with each cell weighted by the local rate at which a given event can happen. The tau-leaping algorithm assumes that $\tau$ is sufficiently small for the number of events to be small in comparison with the number of possible events. This means that the number of events never exceeds the number of available cells and that it is unlikely that two events occur in the same cell at a given time step.
- We update the biotic layers accordingly, and the abiotic layer by calculating all the local changes in resource values within a timestep, then return to the first point for the next time step.

#### 1. 5 Scenarios and settings

Our general focus is to determine the effects of varying dispersal and foraging ranges of species on resource distribution across the landscape. We ran a series of simulations according to a full factorial design crossing the values $\in 1, 2, 5$ (low, medium, high ranges of 9, 25 and 121 cells) for $\Pi_P$, $\Pi_H$, and $\pi_H$, the size of plant and herbivore dispersal ranges, and of the herbivore foraging range, with 25 replicates for each combination. Because these effects likely vary and interact with recycling rates and colonization-extinction dynamics, we consider three scenarios of recycling and eight scenarios of colonization-extinction dynamics. Recycling scenarios comprise an increase of recycling rates from 0, to 1 and 2, keeping the recycling rates of plants and herbivores equal ($\sigma_P = \sigma_H$). Regarding the extinction-colonization dynamics, we considered contrasting scenarios with plant populations displaying either slow $\{e_{P0} = 0.4; \rho_P = 1.6\}$ or fast $\{e_{P0} = 0.9; \rho_P = 3.6\}$ colonization/extinction dynamics, and scenarios with rates that lead to high $\{e_{P0} = 0.4; \rho_P = 2.4\}$ versus low $\{e_{P0} = 1.5; \rho_P = 3.5\}$ plant occupancy. We crossed these scenarios with two scenarios of herbivore colonization-extinction dynamics: fast $\{e_{H0} = 1; \rho_H = 15\}$ and slow $\{e_{H0} = 0.4; \rho_H = 6\}$ dynamics.

The code was implemented in julia language (v1.10). We ran the simulations for cases with plants alone (only varying $\Pi_P$) and with both plant and herbivore populations. We initialized the 100 x 100 cells landscapes with all $N_i$ equal to $N_0$, and all cells occupied by a plant population in all cases. Simulations assume reflecting boundary conditions. To assess how this assumption impacts our results we discarded a 15-cell boundary from the landscapes for the analyses. For scenarios involving herbivores, we randomly set herbivore populations in 20% of the cells. We then ran each simulation during 300 timesteps of the above-described procedure, which was sufficient to reach equilibrium of mean resource values and P and H occupancies in all scenarios. We recorded the final landscapes at the last time point, as well as temporal dynamics of mean resource values and P and H occupancies.

#### 1. 6 Simulation analyses

As outlined in our questions, we mainly focused on understanding effects of foraging and dispersal ranges on resource spatial heterogeneity, and how resource spatial heterogeneity is related to plant and herbivore occupancy and patchiness. To characterize the spatial aggregation in the different layers (resources, plants, herbivores), we calculated different metrics : the mean, variance and skewness of resource levels, the mean of plant and herbivore occupancies, the spatial autocorrelation of resources, plants and herbivores (Moran indices), the join count statistics of plants and herbivores, the three spatial correlations (Tjostheim's coefficients, Tjostheim 1978) between the three layers (pairwise), spectral density ratios (SDR) of the three layers, and the mean, variance and skewness of patch sizes of resources, plants and herbivores (for resources, patch sizes are computed on binarized resource landscapes, obtained through comparison with the median value of resources over the landscape). SDR quantifies the ratio of long-range to short-range correlations (it increases when the relative importance of longer-range correlations increases). Moreover, we calculate the correlations between N-P, P-H and N-H spatial layers to examine the extent to which resource values depended on the presence of plant and herbivore populations. For all statistics that depend on the spatial configuration of the three layers (e.g. Moran indices), statistics are normalized using z-scoring, i.e. removing the expected value obtained from 1000 independent randomizations of resource levels, plant and herbivore occurrences in the final landscape and dividing this centered value by the standard deviation of the same statistic over the 1000 randomized landscapes. Thus, the z-score evaluates the observed spatial structure while controlling for variations in occupancy among scenarios. All analyses were run using R (version 4.5.2.), with package ‘spdep’ (Bivand 2022) for Moran indices and join counts, package ‘SpatialPack’ (Vallejos 2020) for Tjostheim’s spatial correlations, package ‘spatialwarnings’ (version 3.0.3, Génin et al. 2018 MEE) for spectral density ratios and patch sizes.

### Appendix 2: Supplementary figures

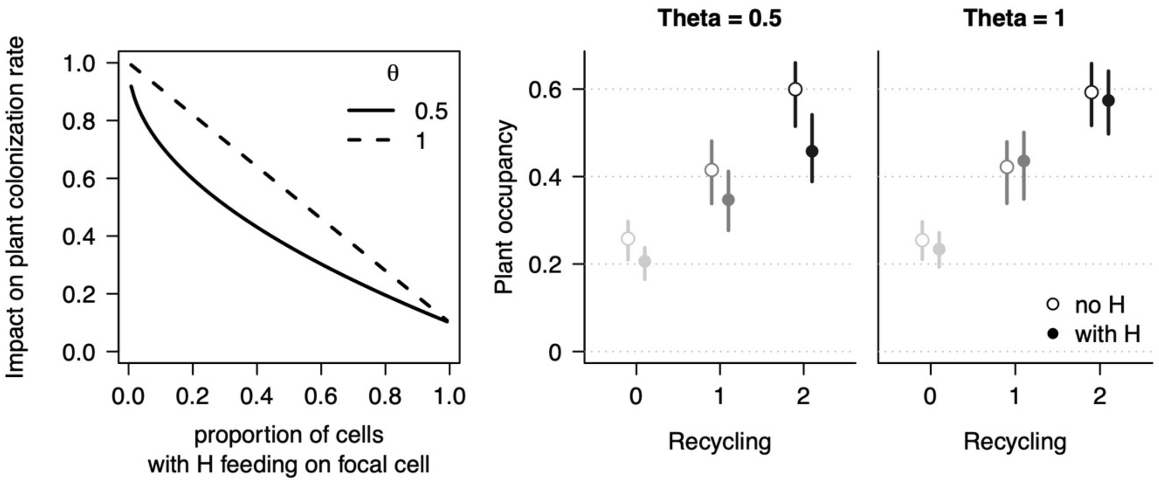

**Figure S1 | Effect of the parameter $\theta$ on plant colonization rate and occupancy.**

Left panel: impact on plant colonization rate in a focal empty cell (multiplicative factor) in function of the proportion of cells occupied by a herbivore in the foraging area surrounding the focal cell for $\theta = 0.5$ (solid line) and $\theta = 1$ (dotted line).

Middle and right panels: Plant mean occupancy in final landscapes without or with the presence of herbivore populations (open and solid circles, respectively) for increasing recycling rates ($\sigma_P = \sigma_H \in \{0,1,2\}$ $with\ corresponding\ gradient\ of\ greys,$ for $\theta = 0.5$ (middle panel) and $\theta = 1$ (right panel). Points and bars give the quartiles. All other scenarios are pooled. See Table 3 for other parameter values. The parameter modulates the shape of the herbivory pressure (see Methods, Appendix 1 and Fig. 1) with smaller values leading to an acceleration of the herbivory pressure on plant colonization rate at low herbivore occupancy (left panels). This generates more top-down dynamics with the presence of herbivores more decreasing plant mean occupancy (middle compared to right panel). The effect increases with recycling level as higher productivity allows higher herbivore occupancy.

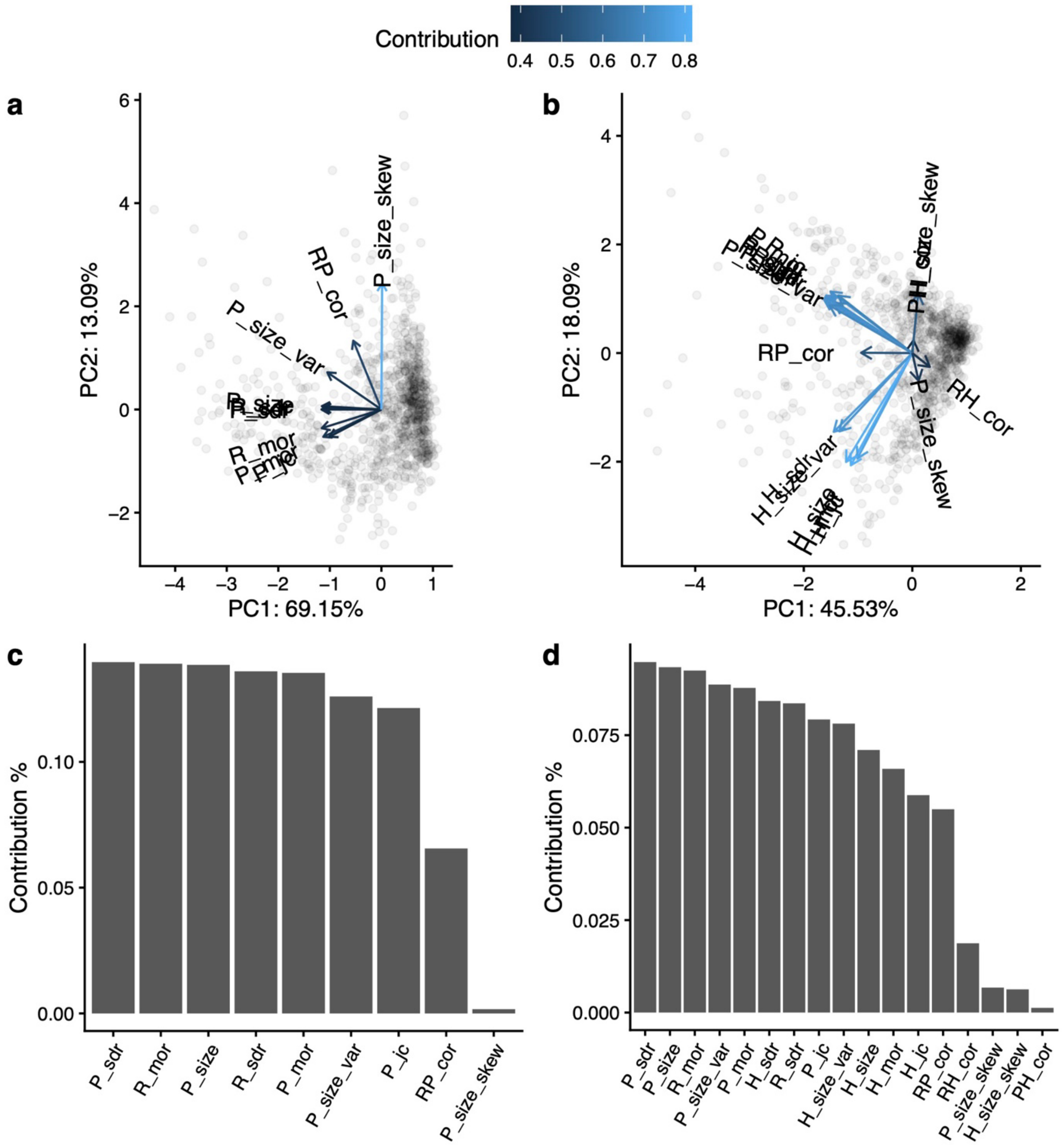

**Figure S2 | Principal Component Analysis on spatial metrics**

for simulations with only plants (a,c) and with plant and herbivore populations (b,d).

a,b) The spatial metrics (arrows) and individual simulations final landscapes (grey circles) projected in the space of the two first axes. c,d) contribution of each spatial metrics to describe the spatial structure of final landscapes. In a and b, this contribution corresponds to arrow colors (see gradient on the top of the panel).

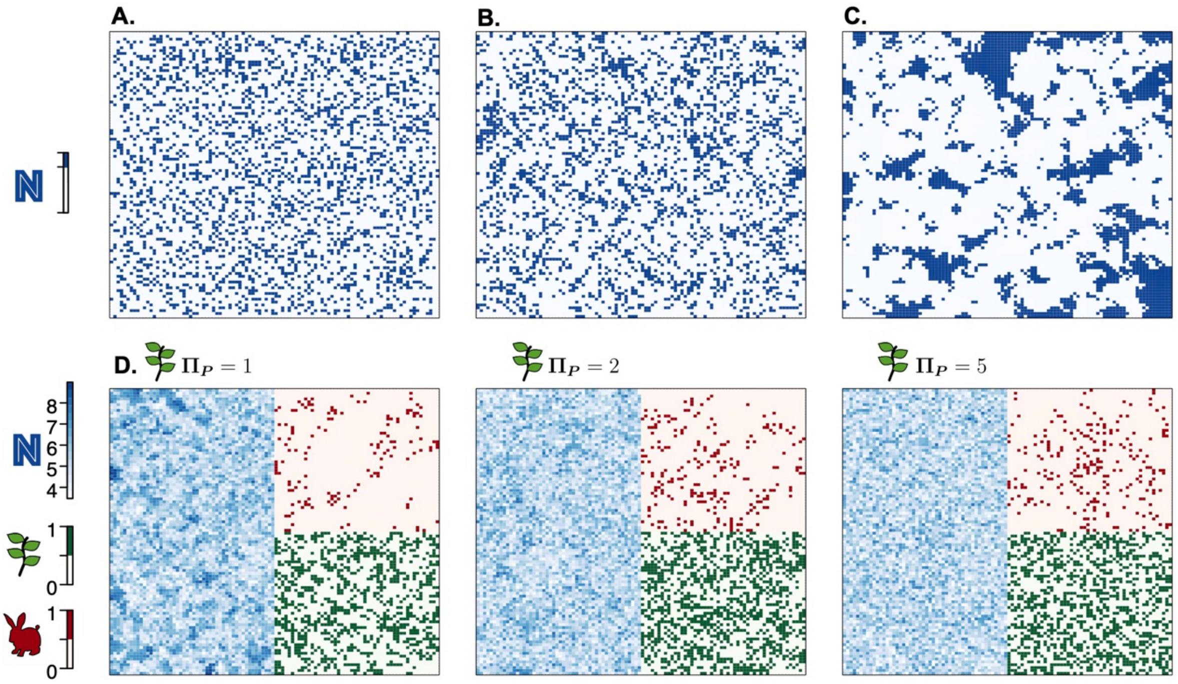

**Figure S3 | Example resource landscapes**

resulting from simulations producing lower-than-expected (A), somewhat higher-than-expected (B), and extremely high (C) spatial structuring of resource distributions (as measured by Moran's I) compared to 1000 randomizations. Blue pixels are those in the highest quartile of resource levels for the given simulation run. Panel (B) represents the median z-score across the whole results dataset, while (A) shows the minimum z-score and (C) shows the maximum z-score. Panel (D) shows distributions of resource levels in the left ⅓ of the landscape, plant occupancy in the middle ⅓ of the landscape, and herbivore occupancy in the right ⅓ of the landscape across three simulation runs that differed only in the parameter value of plant dispersal scale. In panel (D), the z-scores of Moran R for these 3 landscapes are representative of the mean Moran R z-scores of left panels in figure 3.

Parameter values

(A) $\sigma_P = 0.5, \sigma_H = 2, \rho_P = 1.6, e_{P0} = 0.4, \rho_H = 15, e_{H0} = 1, \Pi_P = 5, \Pi_H = 1, \pi_H = 1.$

(B) $\sigma_P = \sigma_H = 1, \rho_P = 1.6, e_{P0} = 0.4, \rho_H = 15, e_{H0} = 1, \Pi_P = 2, \Pi_H = 5, \pi_H = 2.$

(C) $\sigma_P = \sigma_H = 0, \rho_P = 3.5, e_{P0} = 1.5, \rho_H = 15, e_{H0} = 1, \Pi_P = 1, \Pi_H = 2, \pi_H = 5.$

(D) $\sigma_P = \sigma_H = 1, \rho_P = 1.6, e_{P0} = 0.4, \rho_H = 6, e_{H0} = 0.4, \Pi_H = 2, \pi_H = 1.$ $\Pi_P$ variable.

$\theta = 0.5.$ in all panels and see Table 3 for other parameter values.

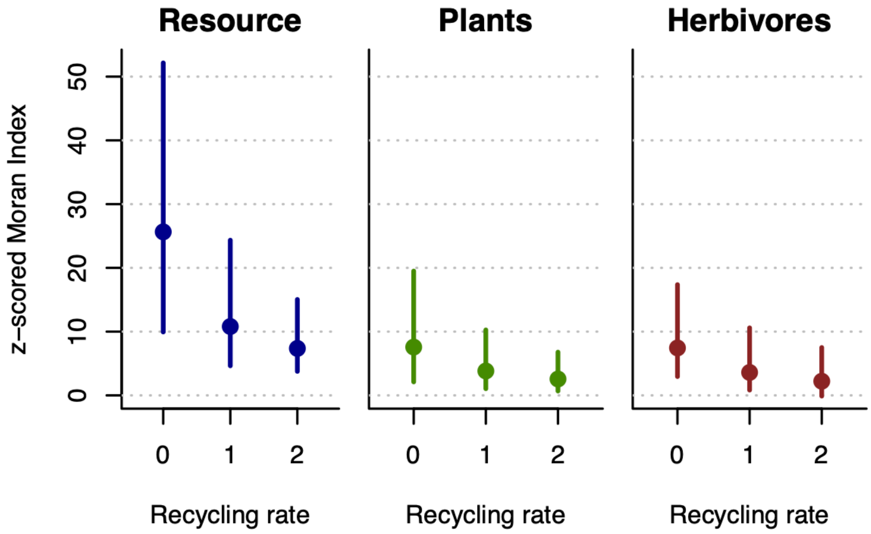

**Figure S4 | Effect of recycling rates on the spatial structure of the three different layers**

(panels and respective colors) as measured by the Z-score of Moran Index of final landscapes (spatial autocorrelation). Moran indices quantify near-neighbor correlation and increases as neighboring cells are more similar and the z-score allows to control for variations in occupancy among scenarios. Points and bars give the quartiles for scenarios with herbivores. All dispersal, foraging ranges and colonization-extinction scenarios are pooled. $\theta = 0.5$. See Table 3 for other parameter values. Raising recycling level has a general negative effect on the spatial structure of all three layers.
